# Single-cell allelic transcriptomic analysis unravels intratumoral gene expression heterogeneity linked to TNBC pathogenesis

**DOI:** 10.1101/2024.04.23.590764

**Authors:** Parichitran Ayyamperumal, Hemant Chandru Naik, Amlan Jyoti Naskar, Srimonta Gayen

## Abstract

Triple-negative breast cancer (TNBC) is the most aggressive subclass of breast cancer, which exhibits high intratumoral heterogeneity. Intratumoral heterogeneity often drives the evolution of drug resistance towards tumour therapy in TNBC. However, the contributing factors and mechanisms behind the intratumoral heterogeneity in TNBC remain poorly understood. It has been implicated that copy number variations (CNVs) often create allelic imbalance and thereby contribute to intratumoral gene expression heterogeneity. However, other processes, such as epigenetic and transcriptional processes, can also contribute to allelic expression heterogeneity in tumour cells. Here, we have investigated how allelic expression heterogeneity contributes to intratumoral heterogeneity in TNBC by performing genome-wide allelic gene expression analysis, excluding most of the CNVs, using single-cell RNA-sequencing datasets. We found widespread monoallelic expression of many genes across different cell types of TNBC. Notably, we found a profound heterogeneity of allelic expression patterns of many genes within TNBC epithelial cells, creating distinct subpopulations of epithelial cells. Importantly, we show that these genes with allelic expression pattern heterogeneity between subpopulations are linked to TNBC immune-surveillance and drug-resistance related pathways, thereby overall shaping the pathogenesis outcomes. Finally, we show that the promoter regions of these genes with profound monoallelic expression are often hypermethylated in TNBC patients. Altogether, our findings reveal that heterogeneous allelic expression patterns contribute to intratumoral heterogeneity and are linked to TNBC pathogenesis.

## Introduction

Cancer is a complex disorder often caused by a plethora of genetic and environmental factors. Emerging studies indicate that cancer cells within a tumour are highly heterogeneous. This intratumoral heterogeneity often results in differential responses towards therapeutic interventions of subpopulations of cells within a tumour (Vitale et al. 2021; Zhang et al. 2016; Kim et al. 2018; Wang et al. 2016)). Triple-negative breast cancer (TNBC) is a highly heterogeneous cancer characterized by loss of estrogen receptor (ER) along with progesterone receptor (PR) and lack of amplification of human epidermal growth factor receptor 2 (HER2) (Dent et al. 2007). TNBC forms 15-20% of all breast cancers and the most aggressive subtypes with poor prognostic options (Bianchini et al. 2022; Reis-Filho and Lakhani 2008). However, the molecular mechanisms driving intratumoral heterogeneity in TNBC remain poorly understood. The recent advent of the single cell RNA-sequencing (scRNA-seq) technology revealed that intra-tumour heterogeneity in TNBC often contributed by copy number variation (CNVs) between subpopulations of cells (Chung et al. 2017; Zhou et al. 2022; Reis-Filho and Lakhani 2008, Karaayvaz et al. 2018). Importantly, it has been demonstrated that CNVs often create allelic expression imbalance and thereby lead to heterogeneous gene expression in cancer (Tuch et al. 2010; Trinh et al. 2022; Rhee et al. 2017; Correia et al. 2022; Chen et al. 2008). However, allelic expression heterogeneity can originate from epigenetic and transcriptional factors as well. Indeed, our studies in the context of development and reprogramming highlighted that heterogenous allelic expression patterns can arise due to the differential transcriptional bursting kinetics of individual alleles of a gene (Ayyamperumal et al. 2024; Naik et al. 2021). Therefore, studying allelic expression heterogeneity based on CNVs only could be insufficient to get the holistic picture. Nevertheless, how the two individual alleles of a gene express to create distinct allelic expression patterns within a TNBC tumour population shapes heterogeneity and subsequent pathogenesis outcomes remains underexplored. Here, we have profiled global allelic gene expression patterns at the single-cell level, excluding most of the CNVs using available scRNA-seq datasets of TNBC patients (Karaayvaz et al. 2018). We found highly heterogeneous allelic expression patterns in TNBC, giving rise to distinct cell populations of epithelial cells within the TNBC tumour. Interestingly, our gene ontology (GO) enrichment analysis revealed that genes with differential allelic expression patterns between cell subpopulations enrich towards cancer immune-surveillance and drug-resistance related pathways. Together, we delineate that heterogeneity in allelic expression patterns might phenotypically shape TNBC pathogenesis outcomes.

## Results

### Widespread monoallelic expression across different cell types of the TNBC tumour

In order to explore the allelic expression patterns at the single cell level of different cell types of the TNBC tumour, we used the already available scRNA-seq datasets from three TNBC patients: PT39, PT84 and PT89 (Karaayvaz et al. 2018). For allelic expression analysis, we performed “variant calling” using the scRNA-seq data to identify the single nucleotide variants (SNVs) between the two alleles, “reference” (Ref) and “alternate” (Alt) for each individual patient. We considered only high-quality SNVs and SNVs, which were not reported in the dbSNP database, were excluded from our analysis. Next, we performed allele-specific analysis following the SNPsplit method with some modifications. First, we profiled the fraction Ref allelic expression of autosomal genes across TNBC single cells of three patients (PT39, PT84 and PT89). As expected, we found that the fraction expression of autosomal genes from Ref allele was ∼0.4-0.6 for most of the cells in all the patients, implicating an overall almost equal gene expression from both of the autosomal alleles (Ref and Alt) (Figure 1A). Next, we profiled SNV-wise allelic expression patterns across different cell types within TNBC tumour, namely, epithelial cells, stroma, macrophage, B cell, endothelial cells and others separately in the three patients. We eliminated most of the SNVs located within CNV regions for individual patients. We observed profound random monoallelic expression of many genes across individual cells of each cell type (Figure 1B). We categorised allele-specific expression patterns into three categories: biallelic, monoallelic expression from the reference allele termed “MonoRef”, and monoallelic expression from the alternate allele termed “MonoAlt”. Altogether, we conclude the presence of widespread random monoallelic expression patterns of genes across different cell types within a TNBC tumour population.

**Figure 1:**
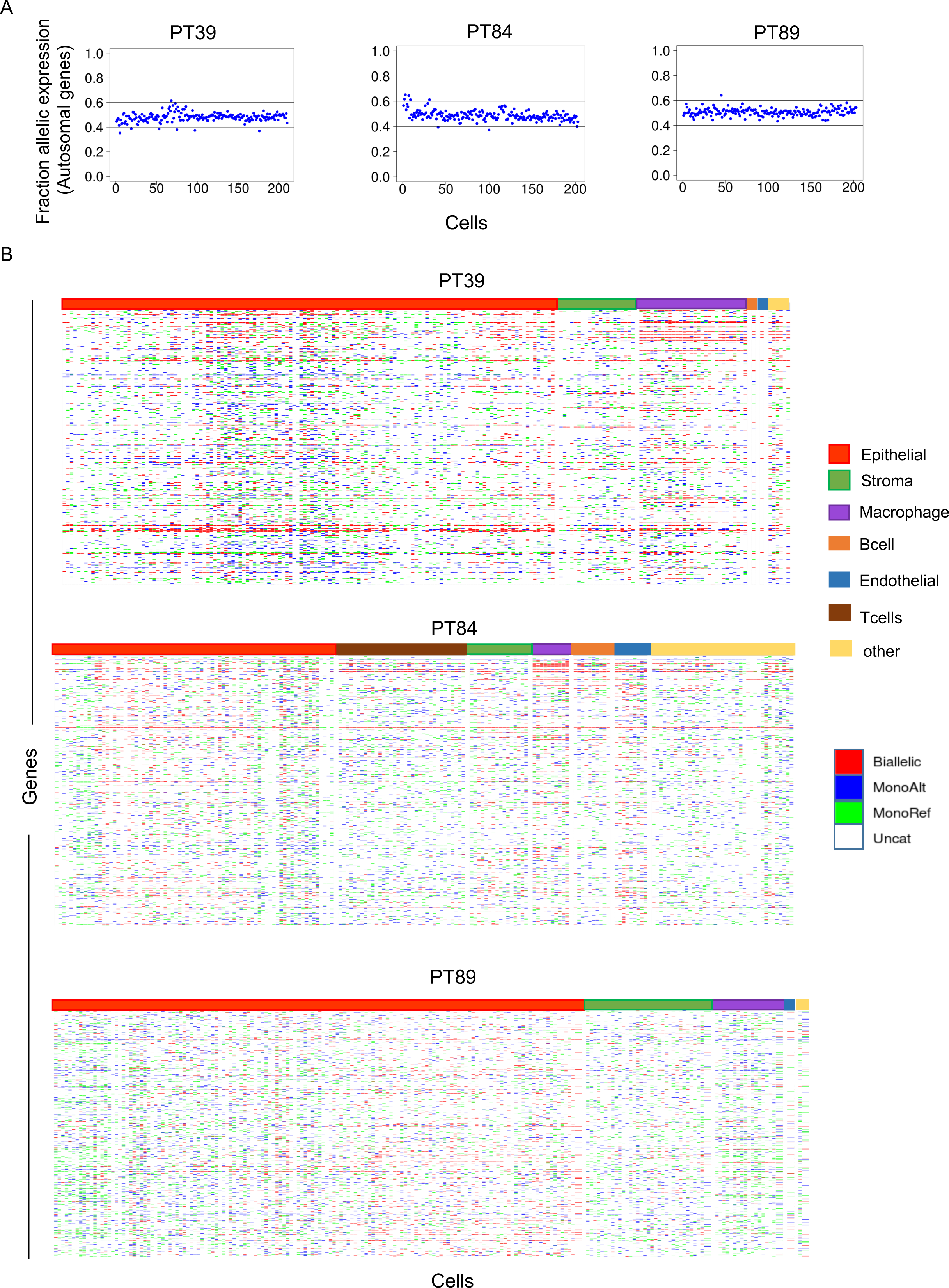
Profiling the allelic expression patterns in TNBC tumour across different cell types. (A) Plot depicting the fraction allelic expression of autosomal genes across TNBC cells in three patients: PT39, PT84 and PT89. (B) Heatmap reflecting the landscape of allelic expression patterns (Biallelic, monoallelic from reference allele, “MonoRef”, monoallelic from alternate allele, “MonoAlt”) across different cell types (epithelial cells, stroma, macrophage, B cells, endothelial cells, T cells and others) in TNBC tumour of three patients, PT39, PT84 and PT89.

### Extensive variation in allelic expression pattern among TNBC epithelial cells

Next, we quantified the percentage of each of the three categories of genes: biallelic, MonoRef and MonoAlt in individual epithelial cells in three TNBC patients (PT39, PT84 and PT89) (Fig. 2). Surprisingly, we found that the majority of the epithelial cells had a very high percentage of monoallelically expressed genes (expressing either from Ref or Alt allele), suggesting widespread monoallelic expression in most of the epithelial cells (Fig 2). On the other hand, we observed that the allelic expression pattern of individual epithelial cells highly varied across the cells in all three patients (Fig. 2). Especially, we found that while some cells harboured a high proportion of biallelic genes, another cohort had an extremely low proportion of genes with biallelic expression across all patients (Fig 2). Based on this, we categorised epithelial cells into three categories: (1) cells that have >60% biallelic genes termed as “high biallelic cells” (2) cells with <10% biallelic genes as “low biallelic cells” and (3) rest of the cells were categorised as “medium biallelic cells” (Fig. 2). We also profiled the percentage of each of the three categories of genes: biallelic, MonoRef and MonoAlt in across other different TNBC cell types such as macrophages, B cell, T cell, stroma etc. in three patients (PT39, PT84 and PT89) (Fig S1). We found that the percentage of biallelic genes in most of the individual cells was low and ranged from 10% to 75% (Fig S1).

**Figure 2:**
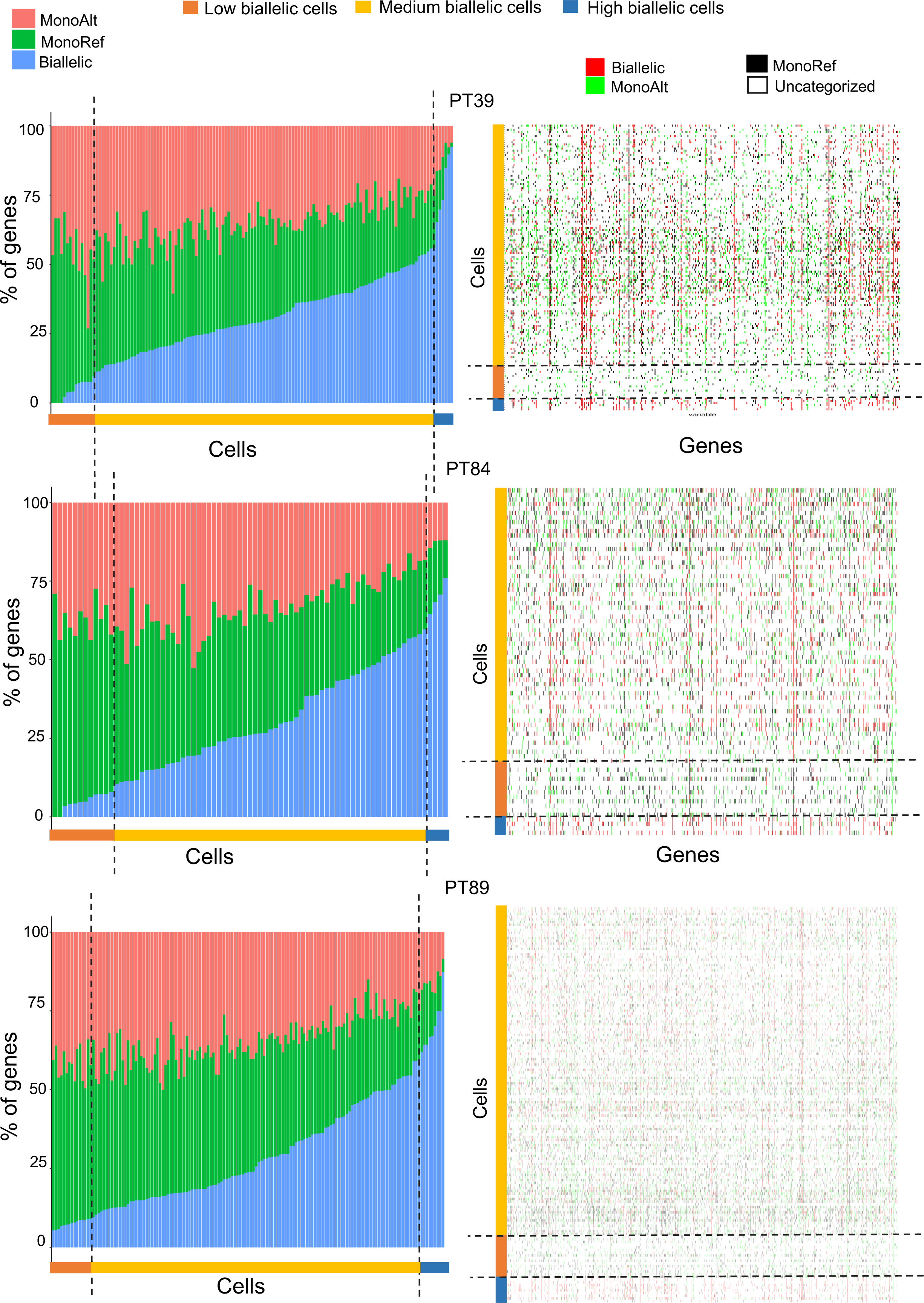
Heterogeneous allelic expression pattern in TNBC epithelial cells. Left: plots representing cell-wise estimation of the percent of the genes with three categories of allelic expression i.e, biallelic, “MonoAlt” and “MonoRef” in low biallelic, medium biallelic and high biallelic epithelial cells (indicated through dotted lines) across three patients, PT39, PT84 and PT89. Right: heatmap reflecting landscape of allelic expression patterns (Biallelic, monoallelic from reference allele, “MonoRef”, monoallelic from alternate allele, “MonoAlt”) in low biallelic, medium biallelic and high biallelic epithelial cells (indicated through dotted lines) across three patients, PT39, PT84 and PT89.

Next, we compared the overall gene expression levels between the high biallelic and low biallelic epithelial cells across all three patients. As expected, we found that in most of the cases, cells of the high biallelic category exhibited higher expression of genes compared to the low biallelic cells (Figure 3). We also compared the gene expression of these two subpopulations with the medium biallelic cells category for all three patients. We found that gene expression was different for high biallelic and low biallelic cells compared to the medium biallelic cells in most of the cases (Fig S2 and S3). Overall, our findings shed light on the relevance of heterogeneous allele-specific gene expression patterns in shaping heterogeneous subpopulations within TNBC tumour epithelial cells.

**Figure 3:**
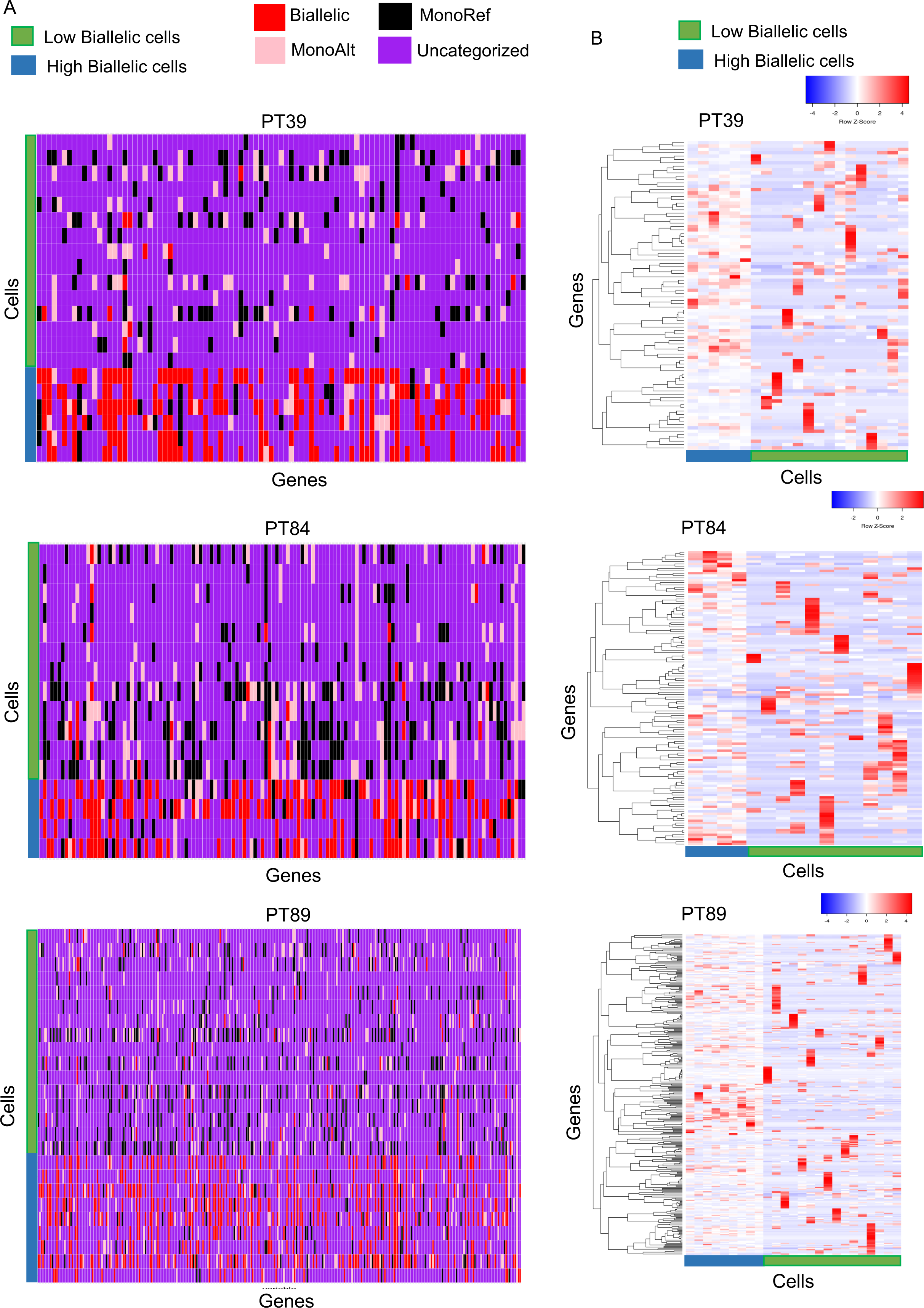
Differential allelic expression pattern between “high biallelic” and “low biallelic” cells correlates with overall gene expression. (A) Heatmap comparing distinct allelic expression patterns between “high biallelic” and “low biallelic” cells in three patients PT39, PT84 and PT89. (B) Heatmap (row Z score of TPM) showing comparison of gene expression levels between “high biallelic” and “low biallelic” cells in three patients.

### Allelic expression heterogeneity is linked to TNBC pathogenesis

In order to gain deeper insights into the pathogenesis outcomes of the allelic expression heterogeneity, we performed gene ontology (GO) enrichment analysis of the genes that showed differential allelic expression patterns between the high biallelic and low biallelic cells (Figure 4). Interestingly, we observed that those genes enriched towards TNBC immune-response related pathways like positive regulation of CD8 positive cells, alpha-beta T-cell proliferation, regulation of T-cell proliferation, positive regulation of T-cell mediated immunity, immune system development, negative regulation of interleukin-2 production, negative regulation of transforming growth factor beta production, leukocyte cell-cell adhesion, cellular response to tumor necrosis factor (Huang et al. 2023a; Li et al. 2018; Huang et al. 2023b; Disis and Stanton 2015; Liu et al. 2020; Vishnubalaji and Alajez 2021; Wakefield and Roberts 2002; King et al. 2004; Fasoulakis et al. 2018) (Figure 4). Furthermore, in PT39 and PT84 patients, some genes enriched towards protection from the natural killer (NK) cell-mediated cytotoxicity pathway (Figure 4). Indeed, NK-cell mediated cytotoxicity is important for anti-metastatic activity in cancer and might have a role in tumour heterogeneity (Yu 2023; Topham and Hewitt 2009). Moreover, in PT39 and PT84 we found genes enriching towards ribosome biogenesis pathways; indeed, ribosome biogenesis is crucial for metastasis and therapeutic resistance in cancer (Metge et al. 2023; Elhamamsy et al. 2022)

**Figure 4:**
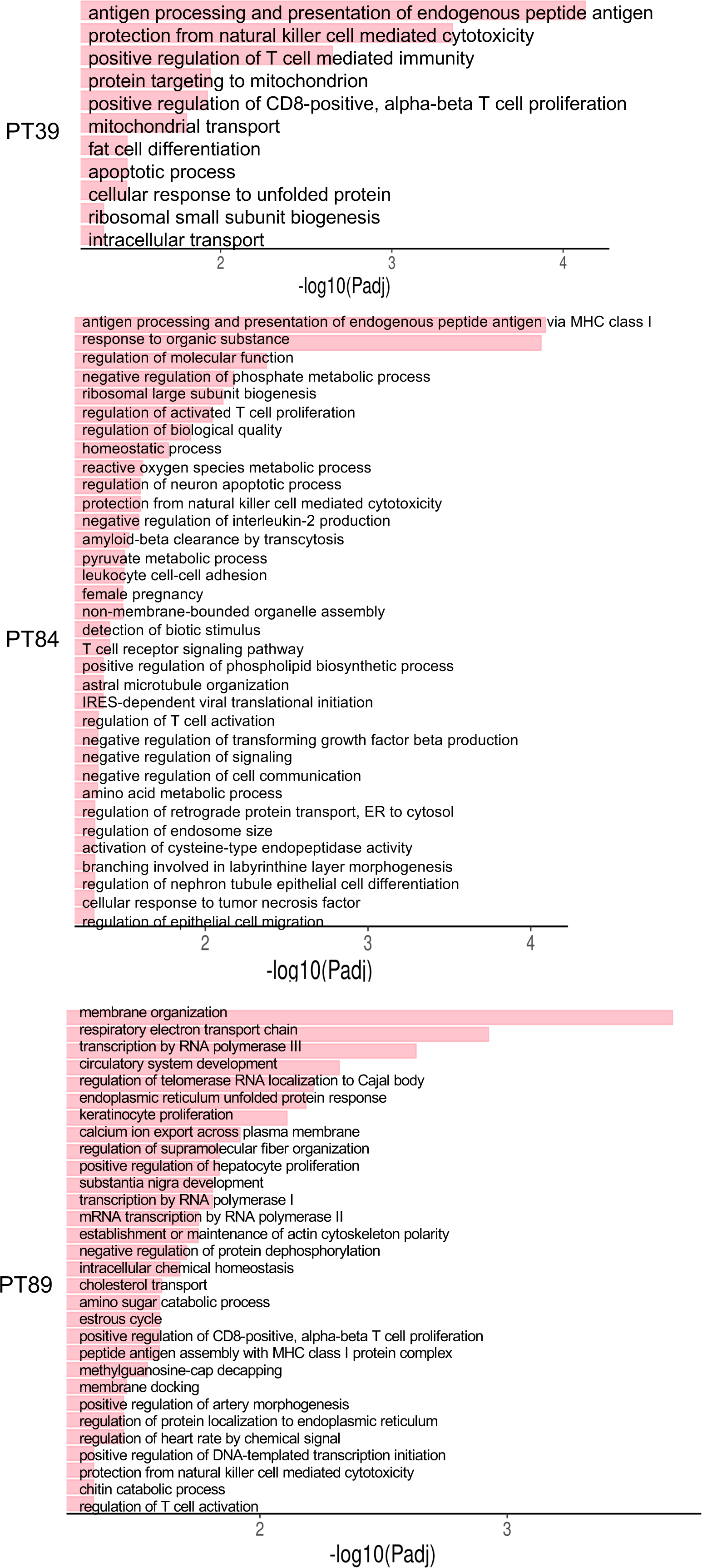
Gene ontology (GO) enrichment analysis. Plots showing GO enrichment analysis of genes with differential allelic expression patterns between “high biallelic” and “low biallelic” epithelial cells in three patients: PT39, PT84 and PT89.

To further explore the link between allelic expression heterogeneity and TNBC pathogenesis, we wanted to determine the prognostic value of the genes that showed differential allelic expression patterns between the high biallelic and low biallelic cell populations. For this, we profiled the association of the expression level of genes enriching towards TNBC immune-response pathways with the survival of TNBC patients (Figure 5). We observed that, in PT39, genes enriching towards cancer immune-response related pathways like positive regulation of CD8-positive alpha-beta T-cell proliferation were associated with relapse-free survival (RFS) of TNBC patients (Figure 5A). Similar results were found for genes enriching in cellular response to tumor necrosis factor, leukocyte cell-cell adhesion, T-cell receptor signaling pathway in PT84 (p < 0.05) (Figure 5A). In PT89, regulation of T-cell activation related genes and peptide antigen assembly with MHC class 1 protein complex further showed similar pattern (p < 0.05). Overall, we demonstrate that the patients with a high level of expression of these immune-related genes have better relapse-free survival (HR= 0.53 – 0.7). On the other hand, we also investigated whether the common genes (TAPBP, HLA-A, CD24 and RAB6A), which showed differential allelic expression patterns across all three patients, are associated with TNBC patients’ survival (Figure 5B). Interestingly, we found that TAPBP is significantly associated with the RFS. High expression of TAPBP in patients is related to better survival (HR=0.6) (Figure 5B). Together, our findings suggested that allelic expression pattern heterogeneity is linked to TNBC pathogenesis and has prognosis values for the survival of TNBC patients.

**Figure 5:**
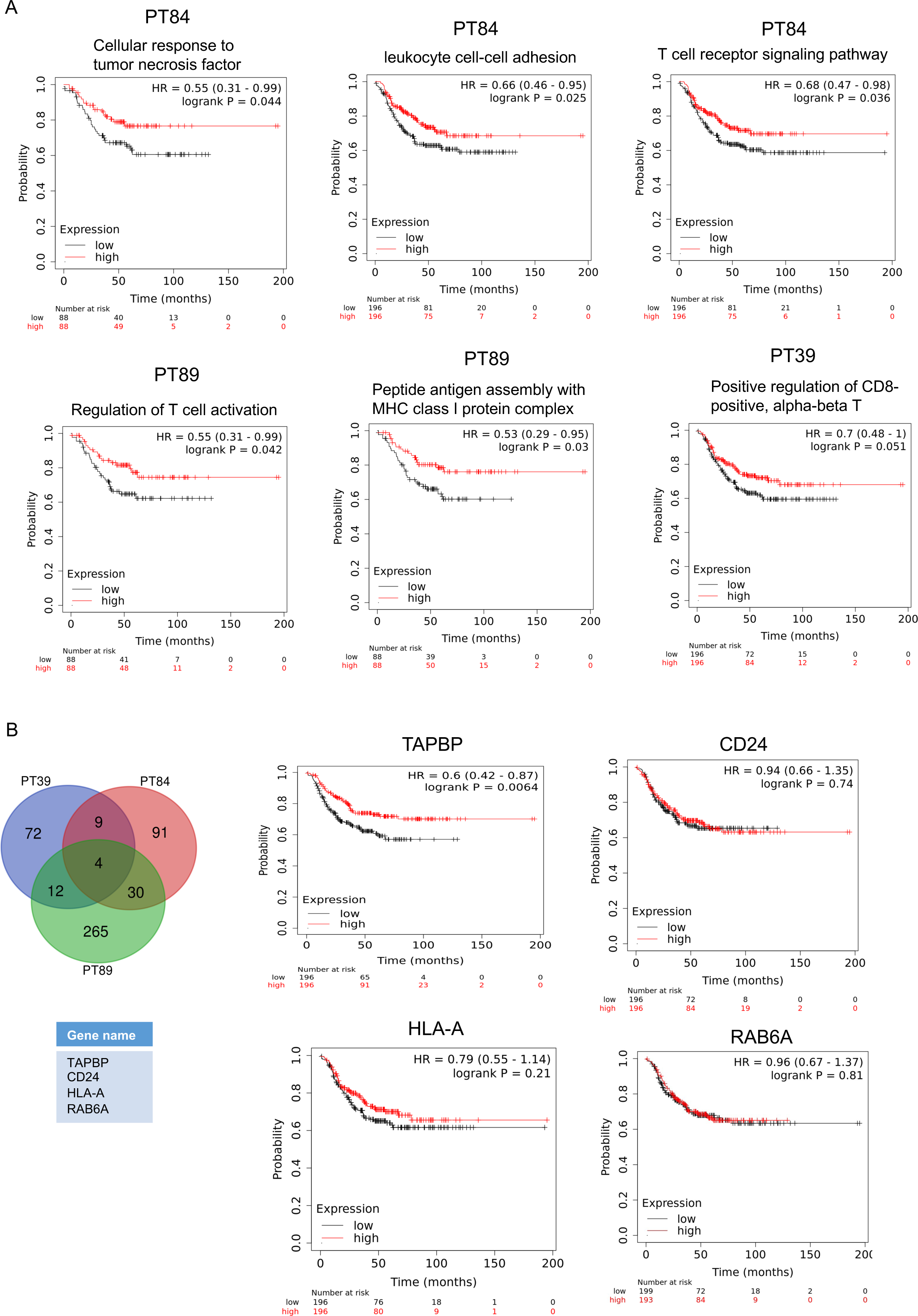
Kaplan–Meier relapse-free survival (RFS) analysis: (A) Plots showing the RFS probability between high and low expression of genes enriched to the significant Biological Process (GO: BP) for each patient. (B)Venn diagram and the table showing four common genes: TAPBP, RAB6A, HLA-A and CD24 across three patients, which showed varied allelic expression patterns between high vs. low biallelic cells. Plots showing the RFS probability between high and low expression of these common genes. Hazard ratio with 95% confidence intervals and log-rank P value were shown.

### Monoallelic gene expression in TNBC is linked to DNA methylation

Next, we explored whether the widespread monoallelic expression patterns within TNBC tumour cells are linked to CpG DNA methylation. Previous studies indicated that DNA methylation is often found to be linked to random monoallelic expression of autosomal genes across different cell types like embryonic stem cells, neural progenitor cells etc. (Marion-Poll et al. 2021; Eckersley-Maslin et al. 2014; Gendrel et al. 2014). Here, we investigated the plausible link between DNA methylation and monoallelic expression within TNBC tumour. For this, we analysed already available MBDCap-Seq datasets to compare the DNA methylation enrichment between normal vs. TNBC cells in the promoter regions of genes that showed differential allelic expression patterns between high biallelic and low biallelic cells (Stirzaker et al. 2015). Interestingly, we observed that promoter regions of the majority of genes are often hypermethylated in TNBC patients (Fig 6A). Moreover, we observed that the promoter regions of four common genes (TAPBP, HLA-A, CD24 and RAB6A), which showed differential allelic expression patterns across all three patients, also showed often hypermethylation in TNBC patients (Fig 6B). Altogether, we conclude that profound monoallelic expression in TNBC cells is often associated with DNA hypermethylation.

**Figure 6:**
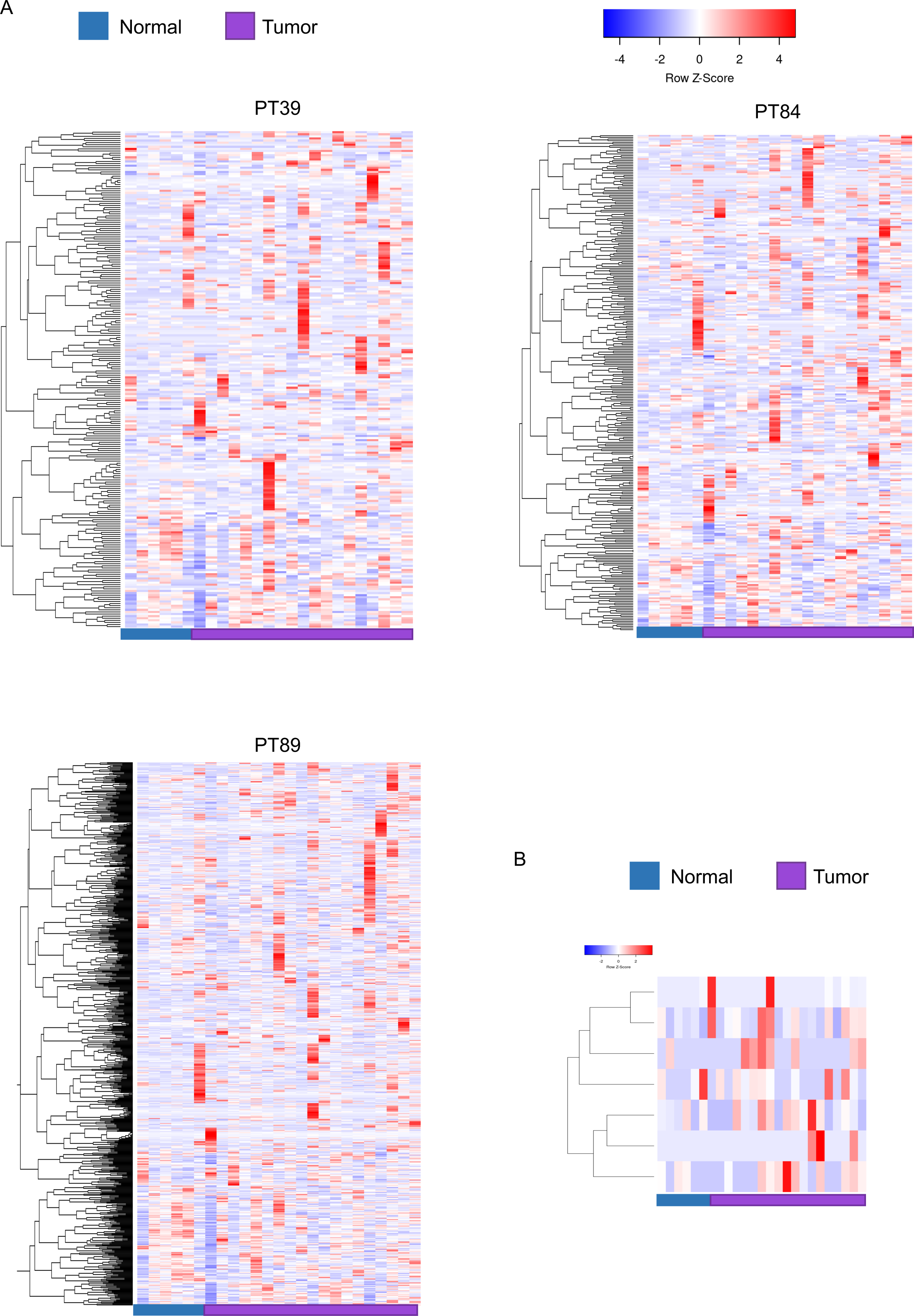
Differential methylation on promoter regions of genes with allelic expression heterogeneity between normal vs tumor. (A) Heatmap showing enrichment of DNA methylation on promoter regions of genes with differential allelic expression patterns between “high biallelic” and “low biallelic” cells in normal vs. tumour cells. (B) Heatmap showing enrichment of DNA methylation on promoter regions of four common genes: TAPBP, RAB6A, HLA-A and CD24 in normal vs. tumour cells.

## Discussion

TNBC is the most aggressive breast cancer subtype in women. TNBC exhibit high intratumoral heterogeneity, which is thought to manifest drug resistance upon treatment. Therefore, understanding the source of intratumoral heterogeneity is imperative for improved prognosis and clinical outcomes (Asleh, Riaz, and Nielsen 2022; Yin et al. 2020; Karaayvaz et al. 2018). In this study, we demonstrate that allelic expression pattern variation in TNBC epithelial cells contributes to intratumoral heterogeneity, which is connected to TNBC pathogenesis, immune-surveillance, drug resistance, etc.

Here, we performed genome-wide allelic expression analyses at the single-cell level using scRNA-seq datasets from three TNBC patients, excluding most of the genes with CNVs (Karaayvaz et al. 2018). Overall, we found widespread random monoallelic expression across different TNBC cell types, like epithelial cells, macrophage, stroma, etc, in all three patients (Figure 1B). As we eliminated most of the CNVs from our analysis, thus this observed random monoallelic expression could have originated largely due to the differential transcriptional burst kinetics between alleles of genes as reported previously (Naik et al. 2021; Ayyamperumal et al. 2024). Nevertheless, we can not completely exclude the possibility of CNV-driven monoallelic expression from our analysis. Indeed, recent single-cell DNA-sequencing (scDNA-seq) suggested that CNVs at the single cell might also result in allelic expression variation and heterogeneity at the single-cell level (Funnell et al. 2022; Shi et al. 2024). In future, better insights can be gained by analysing scDNA-seq datasets to disentangle the role of CNVs vs transcriptional kinetics in shaping allelic expression heterogeneity in TNBC cells.

Next, we profiled the fraction of each category of genes: biallelic, MonoRef, and MonoAlt in individual cells in three TNBC patients. Interestingly, we found two distinctly heterogeneous subpopulations of cells within the epithelial cells: ’high biallelic’ with a higher number of biallelic genes and ’low biallelic’ with a lower number of biallelic genes (Figure 2). Importantly, the allelic expression pattern, as well as overall gene expression, were remarkably different between these two subpopulations (Figures 2 and 3). We further extended our analysis to profile the link between allelic expression heterogeneity in TNBC tumors and pathogenesis outcomes. For this, we performed GO enrichment analysis of those genes that had distinct allelic expression pattern between the ’high biallelic’ and ’low biallelic’ cells. Interestingly, we found many genes enrich towards TNBC immune-surveillance and signalling, and drug-resistance related pathways (Figure 4). Moreover, to gain insights into the prognosis value of allelic expression pattern heterogeneity in TNBC patients, we evaluated the association of expression level of immune-related genes that showed differential allelic expression patterns between “high biallelic” and “low biallelic” cells with survival of TNBC patients. We observed that patients with higher expression of most immune-related pathways have better risk-free survival. On the other hand, we found four genes, CD24, HLA-A, TAPBP, RAB6A to be common among the three patients, changing allelic expression patterns between high biallelic and low biallelic cells that are worth discussing (Figure 5). CD24 has been reported to be responsible for cancer stemness and CD24 expression level determines the prognosis of cancer (Kwon et al. 2015). It is known that patients with higher CD24 expression levels have worse overall survival (OS), distant metastasis-free survival (DMFS) and disease free survival (DFS) than patients with lower levels of expression. CD24 is believed to increase metastatic potential by triggering immune escape. Thus, within the TNBC tumour, cells having higher CD24 expression might have increased metastasis than cells with lower expression (Kwon et al., 2015). On the other hand, previous studies indicated that lesser HLA-A expression levels in TNBC resulted in poor clinical outcomes and it acts as a tumor suppressor (Pujadas and Cordon-Cardo 2021). Moreover, higher level of RAB6A expression is correlated with tumor growth in different cancers (Yang et al. 2023). Furthermore, we performed similar RFS analysis for the genes that commonly switched allelic expression between high biallelic and low biallelic cells. One such gene is TAPBP that was significantly associated with relapse free survival of TNBC patients. Notably, TAPBP is MHC class I antigen presentation related protein that together with TAP1 and HLA-A is responsible for triggering CD8+ T cell activation preventing tumour growth and metastasis in cancer. Hence, indeed, higher TAPBP expression level correlated with better recurrence free survival in TNBC patients (Pedersen et al. 2017).

Separately, we investigated the link between DNA methylation and random monoallelic expression pattern in TNBC. In this regard, our analysis across multiple TNBC tumour samples reflected a higher DNA methylation enrichment of genes in tumour samples than normal samples. This hints about a plausible link between random monoallelic expression pattern and DNA methylation in TNBC. Indeed, differential DNA methylation patterns leading to aberrant gene expression levels have been reported for many cancers (Esteller et al. 2001; Lakshminarasimhan and Liang 2016; Marthong et al. 2020; Batra et al. 2021). Importantly, allele-specific DNA methylation has been found to be linked to certain cancers (Vohra et al. 2021; Qiangwei Zhou et al. 2022). Nevertheless, in future, allele-specific DNA-methylome sequencing analysis can be performed within TNBC to gain better insights into the link between DNA methylation and monoallelic gene expression. Nevertheless, in future, it is imperative to explore the possible interplay among allele-specific DNA methylation, allelic expression patterns and heterogeneity for better prognosis in TNBC. Together, our study sheds light on the allelic expression pattern variation in the TNBC tumour along with its plausible link to epigenetic regulatory mechanisms. Importantly, through this study we shed light on the implications of allelic expression heterogeneity in modulating overall breast cancer heterogeneity along with its pathogenesis outcomes and provides fundamental insights into its prognosis.

## Materials and methods

### Data acquisition

Single-cell RNA-seq datasets of TNBC patients were retrieved from Gene Expression Omnibus (GEO) under the following accession: GSE118390 (Karaayvaz et al. 2018). Methylation sequence data were retrieved from GSE58020 (Stirzaker et al. 2015).

### Identification of single nucleotide variants

To identify single nucleotide variants (SNVs), we performed variant calling using scRNA-seq data of individual patients. In brief, we mapped RNA-seq reads to GRCh38 using STAR 2.710a in individual cells. Next, we merged all the BAM files from single cells for each patient. Then, we performed variant calling using GATK haplotype caller (v4.2.5.0) (Depristo et al. 2011). We removed duplicate reads before variant calling using GATK. We selected high-quality SNVs (QUAL>30) for our analysis. Furthermore, we removed those SNVs which is not present in dbSNP (dbSNP_154).

### Allelic specific RNA-seq analysis

For allele-specific analysis of scRNA-seq dataset, we followed SNPsplit pipeline with some modifications. We removed potential doublet cells from our analysis based on the library size. For allelic expression analysis, we prepared the N-masked genome (GRCh38) using the VCF file generated through the variant calling pipeline as described above. Next, we mapped RNA-seq reads to the N-masked genome and generated allele-wise BAM files using SNPsplit tool (Krueger and Andrews 2016). Then, to get SNV-wise reads, we merged Ref and Alt BAM files and obtained SNV-wise read counts using GATK ASEReadCounter (Depristo et al. 2011). Subsequently, to exclude any false positives, we only considered those SNVs, which had at least 10 reads (Ref+Alt) in an individual cell. Next, we considered those cells for analysis that harboured at least 100 such SNVs. Further, we excluded X-linked SNVs from our analysis. Following this, we selected those SNVs which were expressed in at least 10 cells in individual patient. Also, we excluded those SNVs, which had very biased expression either from Ref or Alt allele across the cells (Ref or Alt allele expressed <20% of cells were removed). Further, we considered 1 SNV/gene. If a gene had more than 1 SNV, we selected a SNV, which had more read depth. We also removed SNVs located within CNV region. CNVs list were obtained from Karaayvaz et al., which they identified based on RobustCNV algorithm using exome sequencing data. We removed duplicated and pseudogenes from our analysis. Finally, we considered only those cells for our analysis, which had at least 10 gene expression. Finally, allelic ratio was calculated individually for each SNVs/gene using the formula = (Ref/Alt reads) ÷ (Ref+Alt reads). A gene was considered monoallelic if at least 95% of the allelic reads came from only one allele.

### Methylome analysis

For profiling DNA methylation, we analysed MBDCap-seq dataset from Stirzaker et al. (Stirzaker et al. 2015). We followed the similar pipeline as described by Stirzaker et al. In brief, we used their processed MBDCap-seq read counts file. We normalized MBDCap-seq data using a fully methylated MBDCap-seq sample dataset (SssI blood sample). We annotated promoter regions with ChIPseeker (G. Yu, Wang, and He 2015). Finally, we visualized the promoter regions of the genes that alter allelic expression between “high biallelic” and “low biallelic” cells in individual patients.

### Gene ontology

Functional enrichment of different classes of genes was profiled using g:GOSt from gProfiler (g:Profiler version e111_eg58_p18_30541362, database updated on 25/01/2024) with Benjamini–Hochberg FDR and selected the biological process having FDR < 0.05 from GO: BP (Kolberg et al. 2023).

### Survival analysis

We used Kaplan–Meier plotter (http://kmplot.com/analysis/) to analyze the RFS (relapse-free survival) in TNBC patients (Győrffy 2021).

## Acknowledgement

This study is supported by Department of Biotechnology (DBT), India (BT/PR30399/BRB/10/1746/2018), Department of Science and technology (DST-SERB), India (CRG/2019/003067), DBT-Ramalingaswamy fellowship (BT/RLF/Re-entry/05/2016) and Infosys Young Investigator award to SG.

## Author contributions

SG conceptualized, supervised and acquired funding for the study. PA performed bioinformatic analyses and contributed to the conceptualization, interpretation of the results and writing of the manuscript. HCN helped with the analysis. AJN wrote the first draft of the manuscript and contributed to conceptualisation. SG, AJN and PA edited the manuscript. The final manuscript was approved by all the authors.

## Declaration

The authors declare no competing interests.

