## Supplementary figures for "Single-cell allelic transcriptomic analysis unravels intratumoral gene expression heterogeneity linked to TNBC pathogenesis"

Fig. S1

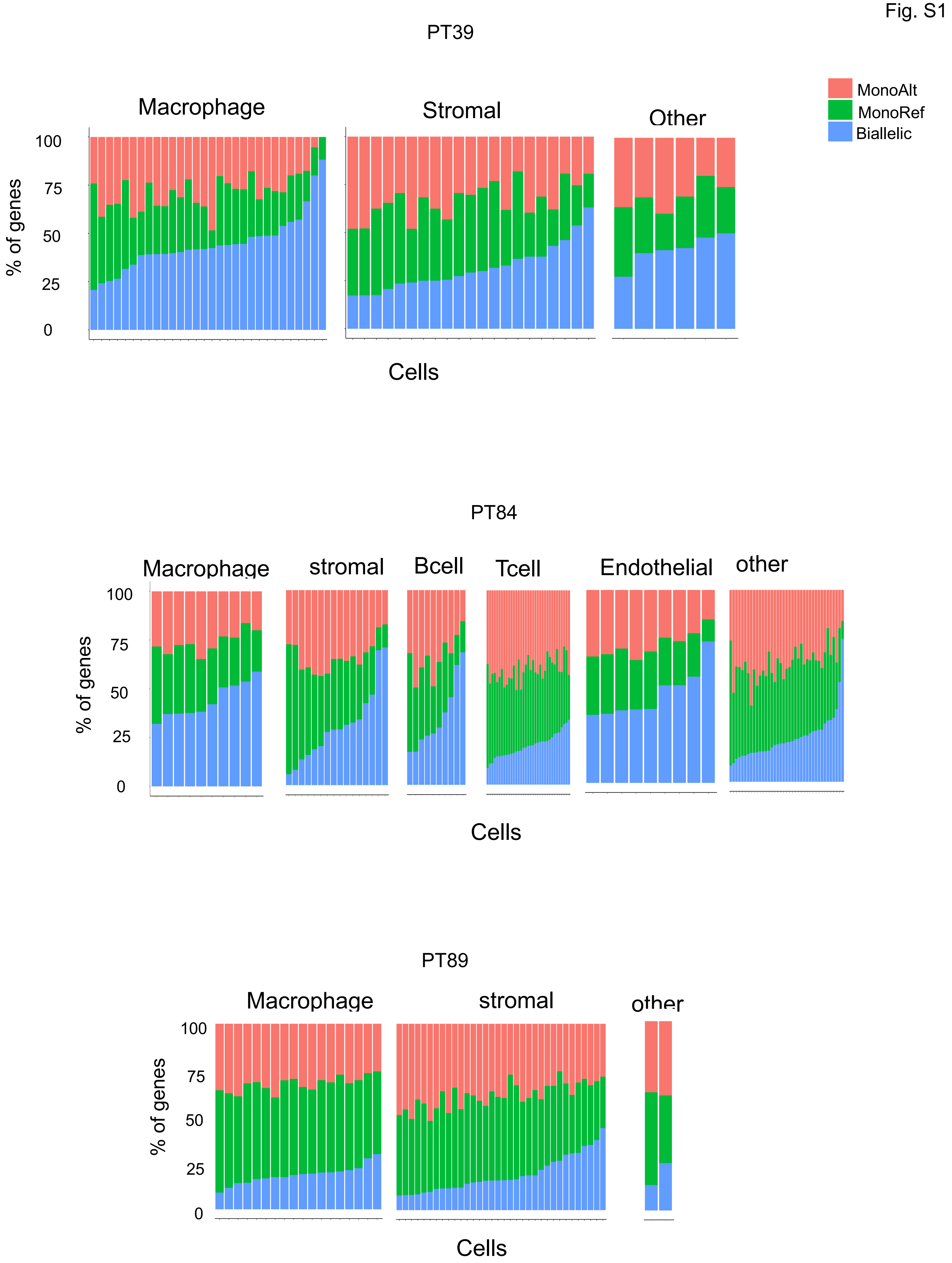
PT39
Macrophage
Stromal
Other
MonoAlt
MonoRef
Biallelic
100
75
50
25
0
% of genes
Cells
PT84
Macrophage
stromal
Bcell
Tcell
Endothelial
other
100
75
50
25
0
% of genes
Cells
PT89
Macrophage
stromal
other
100
75
50
25
0
% of genes
Cells

**Figure S1: Allelic expression patterns across different cell types in TNBC patients:** Bar plots representing the quantification of the percent of three different categories of genes ("MonoAlt", "MonoRef" and "Biallelic") in individual cells within macrophage, stromal and other cells of TNBC tumour in PT39, macrophage, stromal, B cell, T cell, endothelial and other cells in PT84, and macrophage, stromal cells in PT89.

Fig. S2

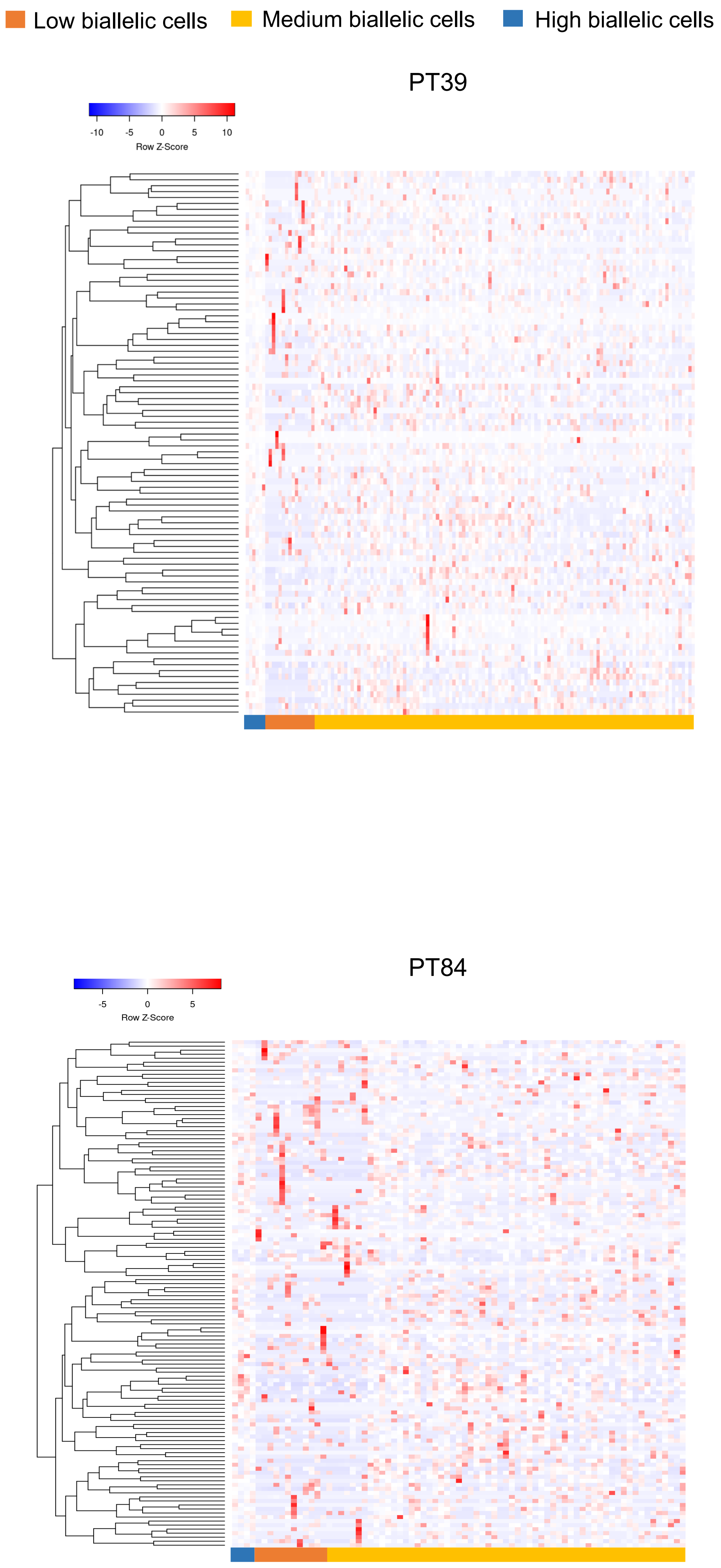
Low biallelic cells
Medium biallelic cells
High biallelic cells
PT39
-10
-5
0
5
10
Row Z-Score
PT84
-5
0
5
Row Z-Score

**Figure S2:** Heatmap illustrating row z-score of TPM of genes with differential allelic expression pattern across three distinct cell populations: high biallelic, medium biallelic and low biallelic cells in patients PT39 and PT84.

Fig. S3

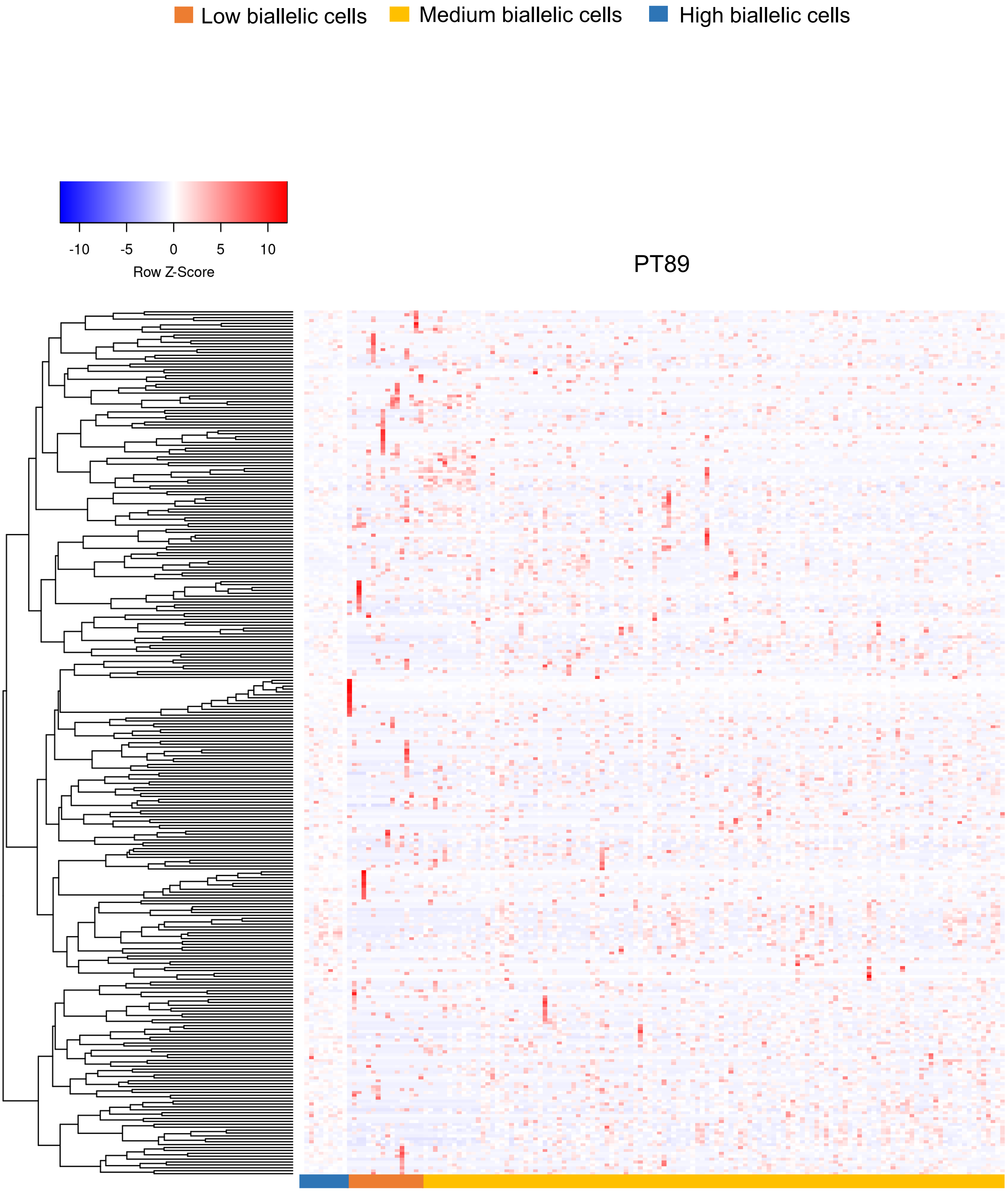
Low biallelic cells
Medium biallelic cells
High biallelic cells
-10
-5
0
5
10
Row Z-Score
PT89

**Figure S3:** Heatmap illustrating row z-score of TPM of genes with differential allelic expression pattern across three distinct cell populations: high biallelic, medium biallelic and low biallelic cells in patients PT39 and PT89.
